# Tau P301S Transgenic Mice Develop Gait and Eye Movement Impairments That Mimic Progressive Supranuclear Palsy

**DOI:** 10.1101/2024.09.20.614197

**Authors:** Rose B. Creed, Scott C. Harris, Sadhana Sridhar, Sascha du Lac, David S. Zee, Felice A. Dunn, Guy Bouvier, Alexandra B. Nelson

**Affiliations:** Kavli Institute for Fundamental Neuroscience, UCSF, San Francisco, CA, 94158; Weill Institute for Neuroscience, UCSF, San Francisco, CA, 94159; Department of Neurology, UCSF, San Francisco, CA, 94158; Department of Ophthalmology, UCSF, San Francisco, CA, 94158; Neuroscience Graduate Program, UCSF, San Francisco, CA, 94158; Department of Otolaryngology-Head and Neck Surgery, Neurology, and Neuroscience, The Johns Hopkins University School of Medicine, Baltimore, MD 21205, USA; Departments of Neurology, Ophthalmology, Otolaryngology-Head and Neck Surgery, and Neuroscience, The Johns Hopkins School of Medicine, Baltimore, USA; Université Paris-Saclay, CNRS, Institut des Neurosciences Paris-Saclay, 91400 Saclay, France

## Abstract

Progressive supranuclear palsy (PSP) is a neurodegenerative disorder with an estimated prevalence of 5-7 people in 100,000. Clinically characterized by impairments in gait, balance, and eye movements, as well as aggregated Tau pathology, PSP leads to death in approximately 5-8 years. No disease-modifying treatments are currently available. The contribution of Tau pathology to the symptoms of patients with PSP is poorly understood, in part due to lack of a rodent model that recapitulates characteristic aspects of PSP. Here, we assessed the hTau.P301S mouse for key clinical features of PSP, finding progressive impairments in balance and gait coordination. Additionally, we found impairments in fast vertical eye movements, one of the most distinctive features of PSP. Across animals, we found that Tau pathology in motor control regions correlated with motor deficits. These findings highlight the utility of the hP301S mouse in modeling key aspects of PSP.

## Introduction

Progressive supranuclear palsy (PSP) is a complex neurodegenerative disorder characterized by cognitive-behavioral and movement symptoms (STEELE et al. 1964). PSP is clinically heterogeneous, but deficits in balance and eye movement are common across most variants and distinguish PSP clinically from other neurodegenerative disorders (Hardwick et al. 2009; Ling 2016; Shaikh et al. 2017). There are no proven symptomatic or disease-modifying therapies for PSP, and the disease leads to death in 5-8 years (Bluett et al. 2021).

At the macroscopic level, several brain regions atrophy in PSP, including cerebral cortex, basal ganglia, midbrain, and brainstem structures (Burciu et al. 2015; Dutt et al. 2016). Microscopically, postmortem studies of PSP found accumulation of the microtubule-associated protein, Tau, in these same regions (Gabor G. Kovacs et al. 2020; STEELE et al. 1964). In the healthy brain, Tau is important in the stabilization of microtubules. Tau is primarily distributed along the axon but can also be found in other neuronal compartments. In PSP, as in other tauopathies, Tau is mislocalized to the dendrites, which may impair its normal functions (Guo et al. 2017). Aggregated Tau is hypothesized to drive the initiation and progression of PSP (Halliday 2000; Robinson et al. 2020). Indeed, recent evidence suggests a causal relationship between pathological Tau and motor impairment: non-human primates inoculated with Tau aggregates extracted from PSP patients developed locomotor impairments (Darricau et al. 2023). However, similar efforts have failed to produce deficits in mice (Narasimhan et al. 2017). Additionally, to date, no study has examined whether a causal relationship exists between Tau pathology and oculomotor impairments in mice. Without a phenotypically penetrant model, investigation of the cellular and circuit mechanisms of PSP have been limited.

Researchers have developed several Tau transgenic mouse lines, which have primarily been used to study the role of Tau in other disorders, such as Alzheimer’s Disease and Frontotemporal Dementia (Denk and Wade-Martins 2009). However, it is unclear how well these mouse lines recapitulate the key clinical features of PSP. One such mouse line, hTau.P301S, overexpresses the predominant Tau isoform found in PSP, 4-repeat (4R) Tau (Allen et al. 2002; Hauw et al. 1994; G. G. Kovacs 2015). These mice have progressive locomotor impairments as well as widespread Tau pathology (Allen et al. 2002; Koivisto et al. 2019; Xu et al. 2014). To determine whether this model recapitulates other key clinical features of PSP, we assessed gait, balance, eye movements, and Tau pathology in hTau.P301S mice. We found age-dependent deficits in coordination and fast vertical eye movements reminiscent of those seen in PSP, suggesting these mice may serve as a model to investigate the pathophysiology of PSP.

## Results

### Gait impairments in hTau.P301S mice

In Progressive supranuclear palsy (PSP), impaired gait and incoordination emerge early (Bluett et al. 2021). To determine whether hTau.P301S mice (hereafter referred to as “Tau mice”) show gait impairments, we performed longitudinal assessments of motor coordination (Fig. 1A, S1A). Consistent with previous reports (Allen et al. 2002), we found hindlimb paralysis at late stages (7+ months), so we examined earlier time points. We found decreased locomotor velocity and impaired performance on the accelerating rotarod in Tau mice at 6 months of age (Fig. S1B; Fig. 1B). Given these robust age-dependent locomotor phenotypes in Tau mice, we next tested whether more subtle changes in gait emerged at even earlier timepoints.

**Figure 1.**
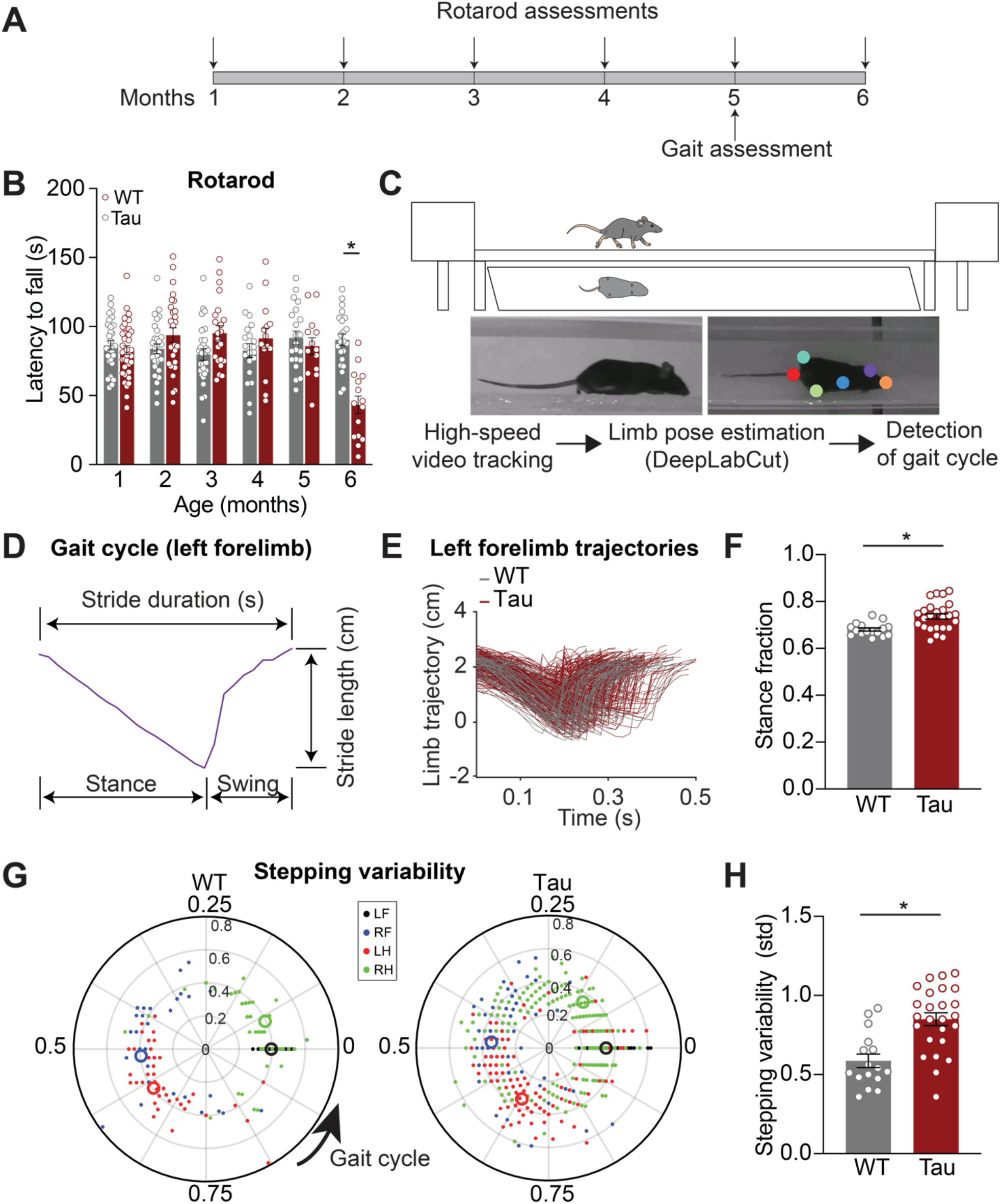
Locomotor coordination decline with age in Tau transgenic mice. **A**. Experimental timeline for locomotor behavior assessment on the accelerating rotarod and linear track in hTau.P301S mice (Tau) and wild-type (WT) controls. **B**. Latency to fall off the accelerating rotarod in WT and Tau mice over the course of 6 months (6 month: N = 22 WT, 15 Tau). *p<0.0001, see Table S3 for full Ns. **C**. Schematic of linear track and mirror (top) used for gait assessment. Limb positions (colored dots) were extracted from high-speed video using DeepLabCut (representative frames, bottom) and processed using custom code to detect the gait cycle. **D**. Example trace of forelimb trajectory during a full gait cycle at 5 months of age. **E**. Overlay of individual forelimb trajectories from one cohort of 5-month-old WT (N = 4, n = 53) and Tau mice (N = 9, n = 259). **F**. Average stance fraction (fraction of time spent in stance phase) in WT and Tau mice (N = 16 WT, 25 Tau). *p = 0.0013.**G**. Polar plots indicating phase of the gait cycle where each limb enters stance, aligned to the stance onset of the left forelimb. Individual dots represent a single stride, different colors correspond to different limbs (LF-left forelimb, RF-right forelimb, LH-left hindlimb, RH-right hindlimb) (representative cohort n = 53 WT, 259 Tau). The distance between the center and the dots represents the duration (s) of that stride. Larger open circles represent the average stride for each limb. **H**. Average stepping variability (standard deviation) across limbs in WT and Tau mice (N = 16 WT, 25 Tau). *p = 0.0002. N refers to mice, n refers to strides. Data are shown as mean +/-SEM. Overlaid open circles in B, F, H represent individual animals.

To determine whether gait abnormalities are present prior to 6 months in Tau mice, we tested 3 - and 5 - month-old mice on a linear track (Fig. 1C, top). We used DeepLabCut, a machine-learning-based method, to track the position of individual limbs during gait (Fig. 1C, bottom) (Mathis et al. 2018). Gait can be divided into individual cycles, each of which includes stance and swing phases, in which the mouse’s paw is on the ground and in the air, respectively. Analysis of these phases yields gait metrics, such as stride length (distance) and duration (time) (Fig. 1D). These gait metrics did not differ between 3-month-old Tau and WT mice (Fig. S1C). However, at 5 months, Tau mice showed several quantitative gait abnormalities, including greater stance fraction (Fig. 1E-F; Table S1). Another element of gait that is impaired in people with PSP is the coordination between body segments; loss of this coordination is associated with frequent falls (Baston et al. 2014). To test whether Tau mice also show this phenomenon, we measured interlimb coordination. In healthy mice, the movement of one limb is closely linked with movement of the other limbs, resulting in tight clustering (low stepping variability, measured as standard deviation) of the phase offset between limbs during the gait cycle (Fig. 1G) (Machado et al. 2015). We found that while 3-month-old Tau mice showed no abnormality in stepping variability, at 5 months they showed greater stepping variability (Fig. S1D-E; Fig. 1H). Together, these findings indicate Tau mice have an age-dependent impairment in gait, balance, and coordination.

### Eye movement impairments in Tau mice

Impaired rapid eye movements are a clinical hallmark of PSP (Kassavetis et al. 2022; Shaikh et al. 2017; STEELE et al. 1964). However, it is unknown whether these impairments can be modeled in Tau mice. To determine whether Tau mice have eye movement deficits, we performed head-fixed eye movement recordings, using video eye-tracking (Fig. 2A). We first tested the integrity of the optokinetic reflex (OKR). The OKR consists of two phases, the first of which is a slow (tracking) phase where the eye follows a moving visual stimulus and, the second of which is a fast, resetting saccade (quick phase of nystagmus). To elicit the slow phase of the OKR, we presented mice with a virtual horizontal oscillating drum, which led to horizontal eye movements that tracked the stimulus (Fig. 2A-B). The drum was rotated at a range of temporal frequencies up from 0.2Hz to 1Hz, eliciting tracking eye movements (Fig. 2C). Both WT and Tau mice showed expected degradation in tracking at higher oscillation frequencies, measured as OKR gain (Fig. 2D) (Van Alphen et al. 2001). These experiments indicate that the slow phase of the OKR is intact in Tau mice.

**Figure 2.**
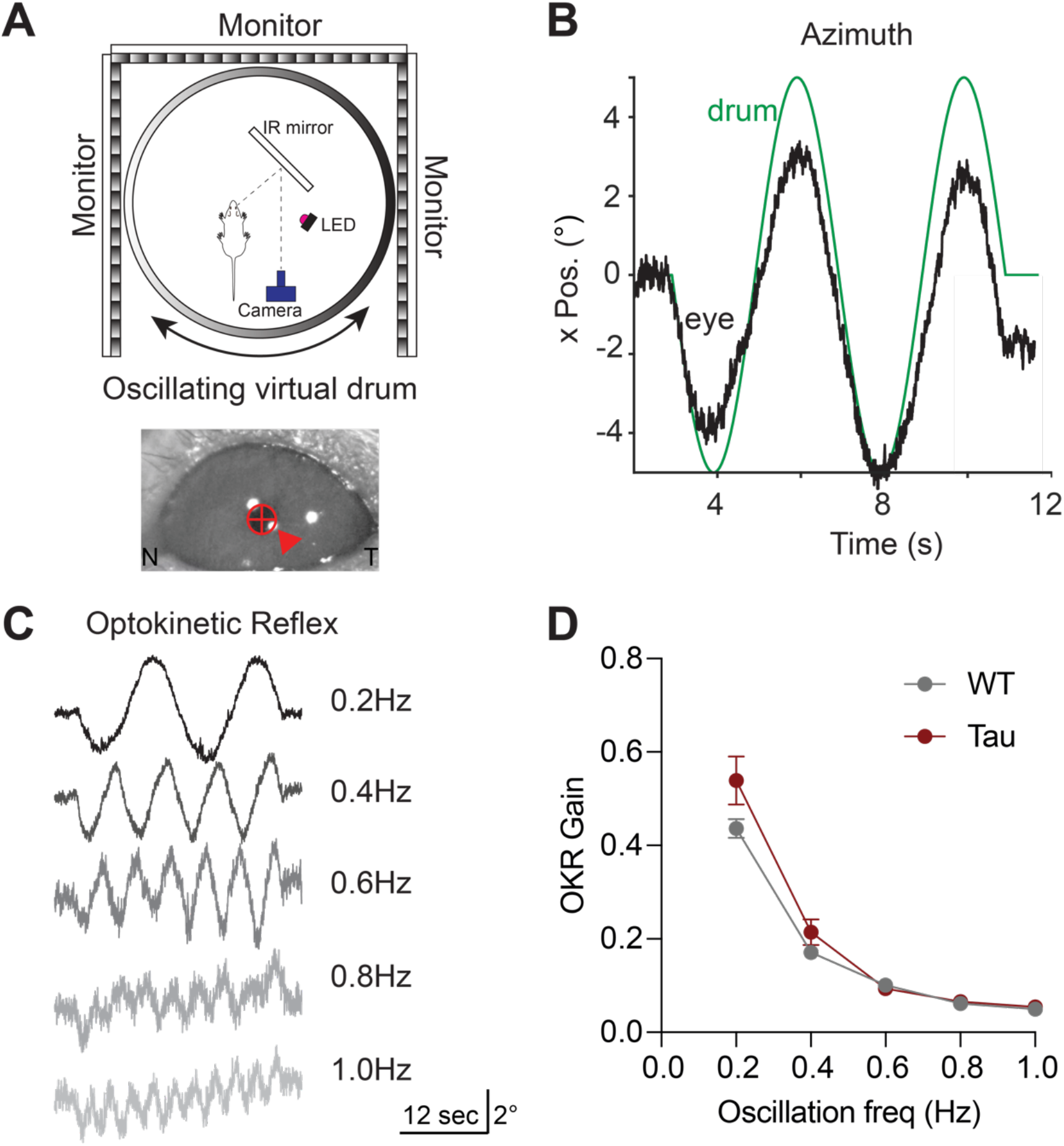
The Optokinetic Reflex (OKR) is Intact in Tau Transgenic Mice. **A**. Schematic of the head-fixed experimental apparatus to measure the OKR (top). OKR was elicited with a virtual rotating striped drum. Image of the mouse pupil as captured during optokinetic stimulation (bottom). Arrow indicates the corneal reflection. N - nasal, T - temporal. **B**. Representative eye movement (black) in the horizontal (azimuth) elicited by sinusoidal rotation of the striped drum (green). **C**. Representative eye movement traces from one mouse in response to optokinetic stimulation at different temporal frequencies. **D**. Analysis of OKR gain (eye movement velocity/drum velocity) in 5-month-old WT and Tau mice (N=9 WT, 7 Tau mice; 0.2 Hz: p = 0.4183, 0.4 Hz: p = 0.6412, 0.6 Hz: p = 0.9955, 0.8 Hz: p = 0.9970, 1.0 Hz: p = 0.9750). Data is shown as mean +/-SEM.

Saccades are rapid, voluntary eye movements that shift gaze to a new location. They complement slower and/or reflexive eye movements that allow tracking of visual stimuli or keep images stable on the retina. In the head-fixed apparatus, we also observed occasional spontaneous (independent of the stimuli) rapid eye movements (>100 deg/sec, >3 deg amplitude) (Fig. 3A). These spontaneous movements had the same kinetic features that are associated with saccades (Fig. 3B-C), and are similar to those reported in recent studies of eye movements in healthy mice (Liu et al. 2016; Sakatani and Isa 2007). We found that these spontaneous eye movements were comparable in WT and Tau mice (Fig. 3D-E; S2A-B). However, in head-fixed mice, the frequency of the spontaneous eye movements is too low to be a practical measure. Moreover, a previous study showed that in people with PSP, resetting saccades (quick phases of nystagmus) are also affected (S. Garbutt et al. 2004).

**Figure 3.**
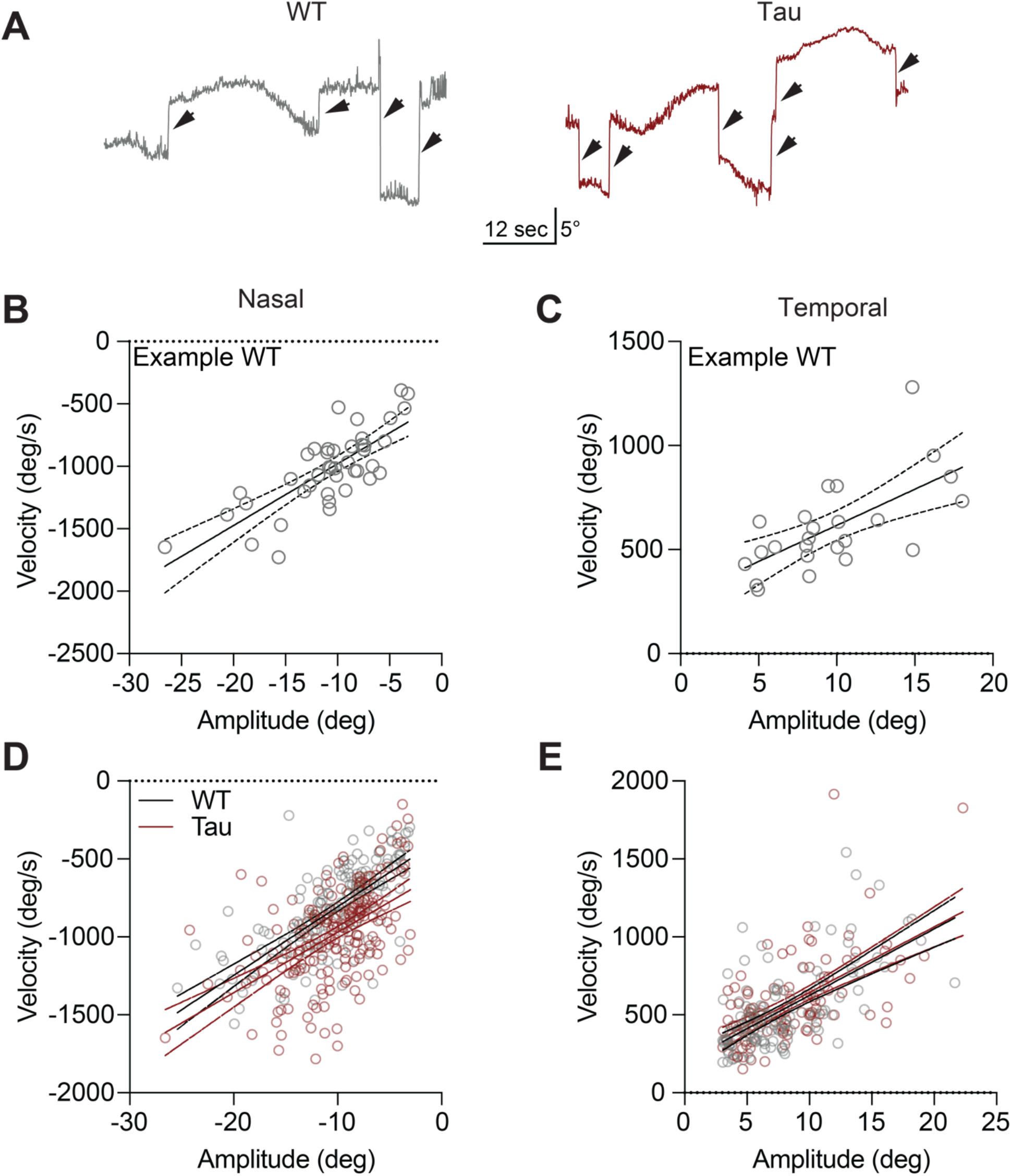
Horizontal saccade-like eye movements are similar in Tau and WT mice. **A**. Representative traces showing rapid (>100 deg/sec) eye movements (arrows) elicited by the horizontally oscillating virtual drum in WT (left) and Tau (right) mice. **B-C**. Representative peak velocity-amplitude relationship of (B) nasal and (C) temporal eye movements in a WT mouse. **D-E**. The velocity-amplitude relationship of nasal (D) and temporal (E) eye movements in the entire cohort (N = 8 WT, 6 Tau mice). Each circle represents a single saccade-like eye movement. Solid line = linear regression, dashed lines represent 95% CI.

To more sensitively test the integrity of both horizontal and vertical saccade-like eye movements, and to evoke more of these eye movements, we measured resetting saccades during visual tracking. Resetting saccades are particularly prominent when using rapid, continuous visual stimuli, rather than slow, oscillating stimuli. To most efficiently elicit resetting saccades, mice were headfixed inside a dome, allowing for full coverage of the visual field, and presented with unidirectional stimuli moving at the equivalent of 10 deg/sec (Fig. 4A). Mice tracked stimuli in temporal, nasal, ventral, and dorsal directions (Fig. 4B-C), showing similar OKR slow-phase gains in Tau and WT mice (Fig. 4D). We then quantified the kinematics of the resetting saccades. Horizontal resetting saccades were observed with temporal and nasal stimuli, showing comparable amplitude-peak velocity relationships in WT and Tau mice (Fig 4E-F; Fig. S2C-D).

**Figure 4.**
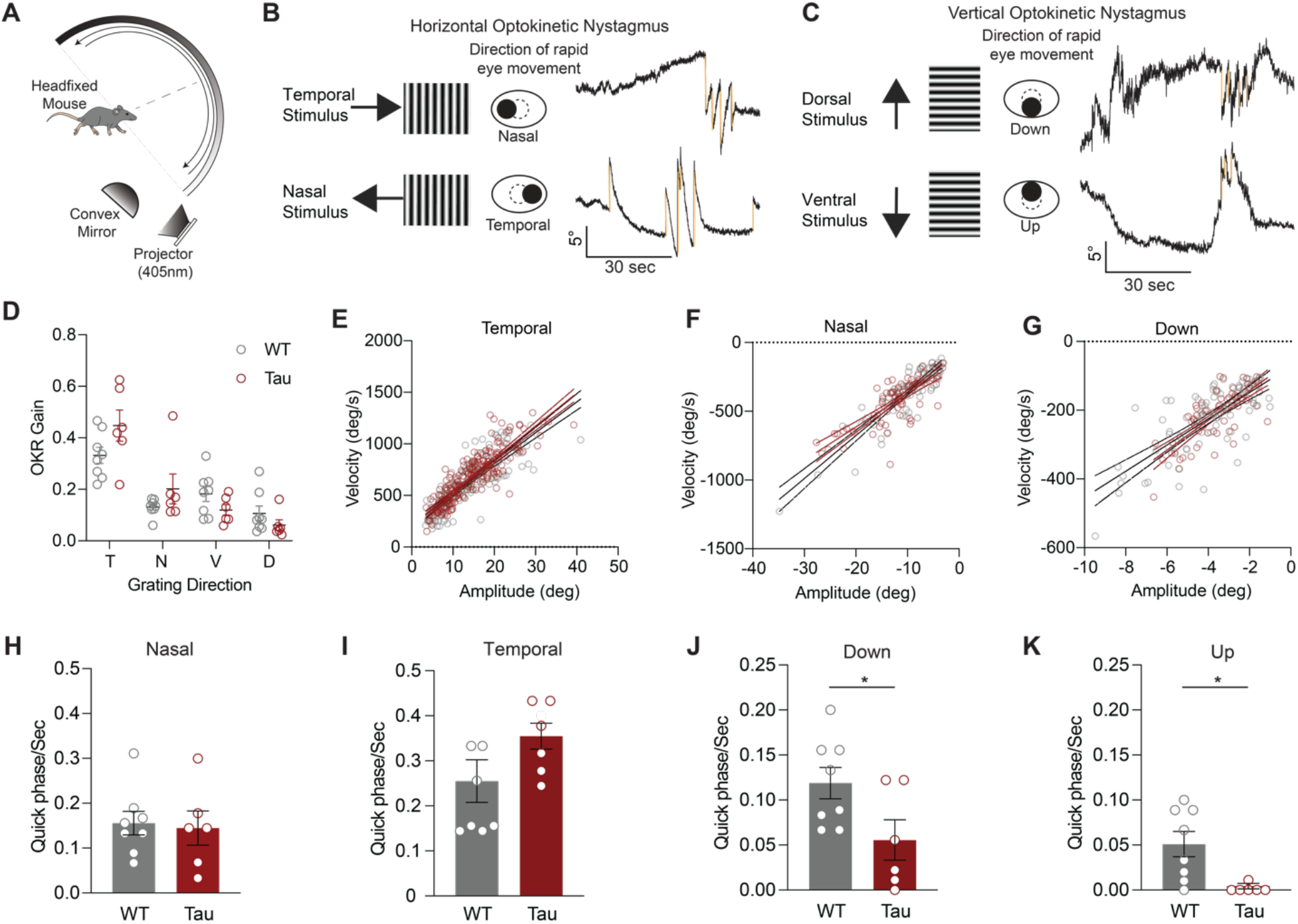
Tau Mice Make Fewer Vertical Resetting Saccades. **A**. Head-fixed apparatus to measure eye movements in response to continuous, full-field visual stimuli. Each mouse was placed in the center of the hemisphere and positioned to record movements of the left eye with a high-speed camera. An optokinetic stimulus was projected onto a mirror and reflected on the hemisphere. Stimuli were presented horizontally or vertically in different trials. **B**. Assessment of the <u>horizontal</u> optokinetic response. As the grating moved in the temporal direction (top), mice made slow, tracking movements in the temporal direction (‘slow phase’), and rapid eye movements (‘quick phases’) in the nasal direction to reset. Quick phases, or resetting saccades, are shown as downward deflections (gold) in eye position. Similar analysis can be performed with the grating moving in the nasal direction (bottom). **C**. Assessment of <u>vertical</u> optokinetic response. Top: as the grating moved in the dorsal (upward) direction, mice made upward slow tracking movements, and more rapid ventral (downward) resetting saccades. These are shown as downward deflections in eye position (gold). Bottom: as the grating moved in the ventral (downward) direction, mice made downward slow tracking movements, and more rapid upward resetting saccades. These are shown as upward deflections in eye position. **D**. OKR (slow phase) gain in the temporal (T), nasal (N), ventral (V), dorsal (D) stimulus directions in WT and Tau mice (N = 8 WT, 6 Tau. T: p = 0.5185, N: p > 0.9999, V: p = 0.4430, D: p = 0.8834). **E-G**. Main sequence (peak velocity-amplitude relationship) of resetting saccades in the (E) temporal, (F) nasal, and (G) downward directions in WT and Tau mice. Mice made too few upward resetting saccades to be displayed in this fashion. **H-I**. Frequency of horizontal quick phases in the nasal (H) and temporal (I) directions WT and Tau mice (N = 8 WT, 6 Tau. D, p = 0.8248. E, p = 0.1678). **J-K**. Quantification of the frequency of ventral (J) and dorsal (K) quick phases in WT and Tau mice (N = 8 WT, 6 Tau; p = 0.0373; p = 0.0077). Data is shown as mean +/-SEM. N = mice.

In people with PSP, the velocity, amplitude, and frequency of vertical saccades are preferentially impaired (Chen et al. 2010). To test whether Tau mice show similar pattern of deficits, we analyzed vertical resetting saccades. We found that the frequency and amplitude of vertical resetting saccades was lower than horizontal resetting saccades, consistent with prior studies of the vertical OKR (Al-Khindi et al. 2022; Yonehara et al. 2016). For this reason, we could only assess the amplitude-peak velocity relationship of downward resetting saccades; these were similar across genotypes (Fig. 4G; Fig. S2E). The frequency of horizontal resetting saccades (in either direction) was comparable between WT and Tau mice. (Fig. 4H-I). However, Tau mice made fewer vertical resetting saccades than WT mice when provided the same stimuli (Fig. 4J-K, supplementary Movie 1 and 2). Indeed, Tau mice had almost zero upward resetting saccades (Fig. 4K). These results indicate that Tau mice have a lower frequency of vertical resetting saccades, which has also been observed in people with PSP (S. Garbutt et al. 2004).

### Tau aggregation correlates with behavior impairments in Tau mice

Tau aggregates are the neuropathological hallmark of PSP. We next examined where pathological Tau accumulates in Tau mice, and how this correlated with motor deficits. Tau mice develop pathology in many brain areas, but prior quantitative histology focused on the hippocampus (Koller et al. 2019). Here, we expanded this analysis to brain regions relevant for motor control, many of which show early Tau pathology in PSP itself (Gabor G. Kovacs et al. 2020). We immunostained brain sections from Tau mice for phosphorylated Tau, a validated marker of pathological human Tau (Goedert et al. 1995). Consistent with previous reports (Allen et al. 2002; Koivisto et al. 2019), we detected phosphorylated Tau immunoreactivity in many brain regions in 5–6-month-old Tau mice (Fig. 5A). By comparison, phosphorylated Tau was absent in age-matched WT controls and minimal in 3-month-old Tau mice (Fig. S3A-B). In 5–6-month-old Tau mice, pathological Tau was found in several motor control regions, including the motor cortex (MCtx), superior colliculus, motor region (SCm), zona incerta (ZI), substantia nigra pars reticulata (SNr), pedunculopontine nucleus (PPN), cuneiform nucleus (CUN), subthalamic nucleus (STN), vestibular nucleus (VN), and deep cerebellar nuclei (DCN) (Fig. 5B). These findings extend prior descriptions of pathology in Tau mice to premotor nuclei within subcortical circuits controlling movement, including many that are associated with eye movement.

**Figure 5.**
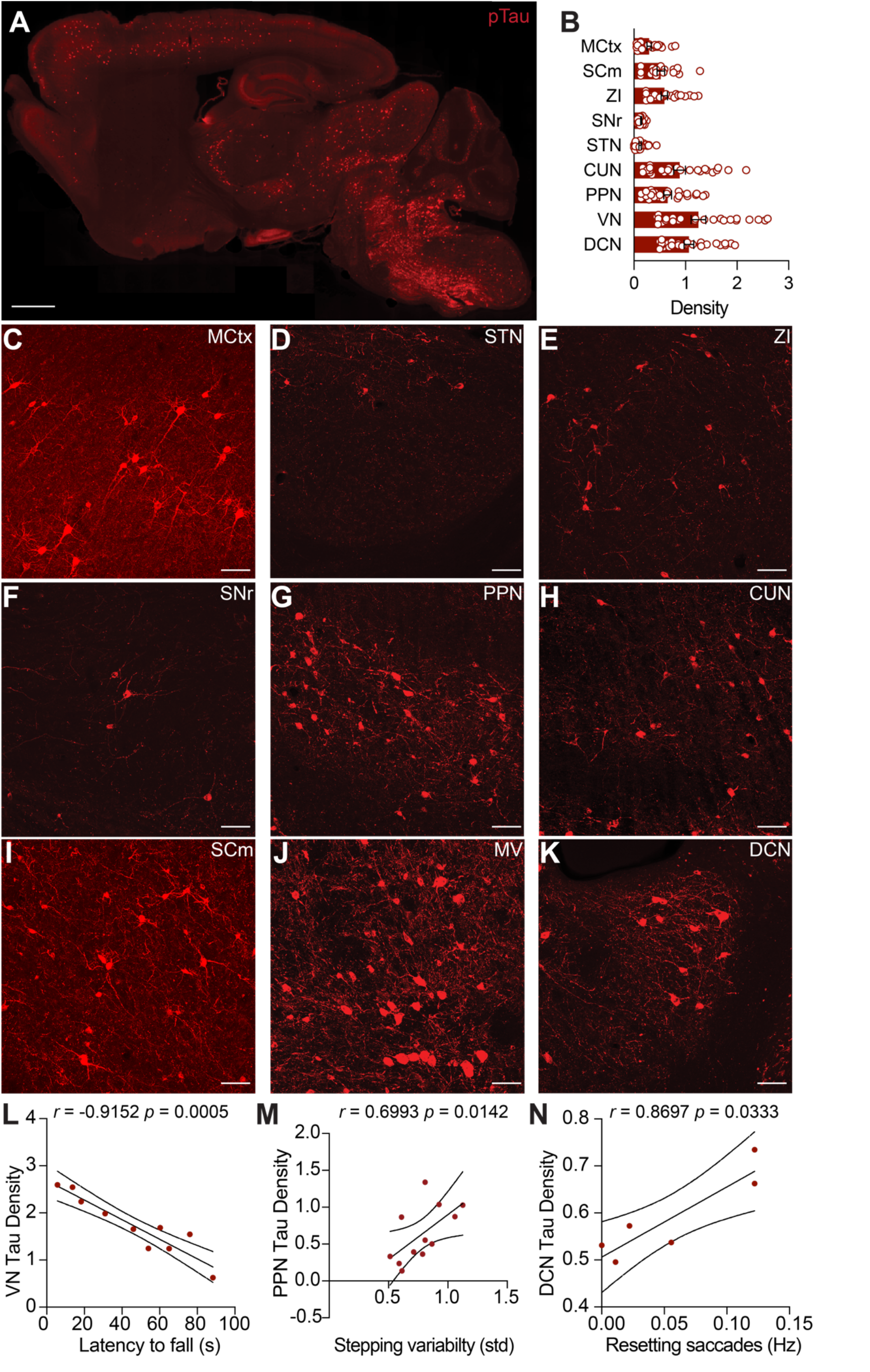
Motor Impairments Correlate with Density of Tau Pathology in Subcortical Areas. **A**. Representative sagittal section from a 5-month-old Tau mouse immunostained for phosphorylated Tau (AT8 antibody). Immunoreactivity was detected in many areas of the brain. Scale bar = 1mm **B**. Summary of density (AT8+ cell counts/10^4^ µm^2^) of Tau pathology in key motor control regions in 5–6-month-old Tau mice. Regions include the Motor cortex (MCtx), Superior colliculus, medial (SCm), Zona incerta (ZI), Substantia nigra pars reticulata (SNr), Subthalamic nucleus (STN), Cuneiform nucleus (CUN), Pedunculopontine nucleus (PPN), Vestibular nucleus (VN), and the Deep Cerebellar Nuclei (DCN). Overlaid circles represent individual mice. N = 26 mice **C-K**. Representative high magnification images showing Tau pathology in the (C) MCtx, (D) STN, (E) ZI, (F) SNr, (G) PPN, (H) CUN, (I) VN, (J) DCN, (K) SCm. Scale bars = 50µm **L**. Correlation between the density of Tau pathology in the VN and performance on accelerating rotarod at 6 months. **M**. Correlation between density of Tau pathology in the PPN and stepping variability on the linear track at 5 months. **N**. Correlation between density of Tau pathology in the DCN and downward quick phase frequency at 5 months. Significant correlations and their coefficients are indicated on each panel. See Table 2 for correlations between pathology and motor impairment across all areas examined. Filled symbols represent individual mice. N = 10 mice for latency to fall, 12 mice for stepping variability, 6 mice for quick phases.

We observed variability among animals in both behavior and pathological Tau density. To test whether these observations might be related, we looked for correlations between behavioral measures (performance on the accelerating rotarod, stepping variability, or vertical resetting saccades) and the density of Tau pathology (Fig. 5C-K; Table S2). Amongst the regions we examined, we found pathology in the VN, an area important in integrating vestibular input, correlated with rotarod performance (Fig. 5L; Table S2). Moreover, pathology density in the PPN, DCN, and zona incerta (ZI) negatively correlated with rotarod performance, and positively correlated with stepping variability on the linear track (Table S2, 5M). Lastly, we tested whether the frequency of vertical resetting saccades correlated with Tau pathology density. Amongst the brain regions examined, we found that DCN pathology correlated with vertical saccades (Fig. 5N, Table S2). These results indicate that Tau pathology accumulates in subcortical motor circuitry in parallel with the development of locomotor and oculomotor impairments in Tau mice.

## Discussion

The main finding of this study is that hTau.P301S (Tau) transgenic mice recapitulate several distinctive clinical features of PSP: deficits in gait, balance, and saccadic eye movements. The gait impairments preceded the onset of gross coordination deficits. We found the slow phase of the optokinetic reflex (OKR) is preserved in Tau mice. However, Tau mice showed selective impairment in vertical resetting saccadic eye movements (quick phases), much as in early PSP. These deficits correlate with the accumulation of pathological Tau in key subcortical nuclei. These observations not only expand our understanding of locomotor impairments previously reported in Tau mice, but for the first time identify ocular motor impairments pathognomonic of PSP.

Our findings add to a body of evidence indicating that Tau transgenic animals develop balance and gait impairments reminiscent of those seen in PSP (Jang et al. 2019). As in previous studies in hTau.P301S mice, we found progressive locomotor impairments (Allen et al. 2002; Koivisto et al. 2019). However, the timing of these impairments may be strain-specific: in our C57Bl/6 background mice, deficits appeared at 5 months of age, similar to another study on this background (Xu et al. 2014), but somewhat later than seen in previous studies of Tau mice on a mixed CBA/C57BJ6 background (Allen et al. 2002; Koivisto et al. 2019). More importantly, we demonstrated that Tau mice showed quantitative gait deficits that parallel those seen in people with PSP (De Vos et al. 2020; Egerton et al. 2012).

Deficits in saccadic eye movement are a distinctive, early and often diagnostic feature of PSP. Until later stages, people with PSP have intact OKR (Chen et al. 2010; Joshi et al. 2010). However, they have impairment in saccadic eye movements, with vertical saccades typically affected before horizontal saccades (Shaikh et al. 2017). Until now, eye movements have not been studied in Tau transgenic mice. To test whether Tau transgenic mice might be a viable model for PSP, we assessed several types of eye movements in Tau mice. We found that WT and Tau mice tracked visual stimuli in both horizontal and vertical directions, with comparable OKR gain. This observation mirrors early PSP (Chen et al. 2010).

We next examined saccadic eye movements. There are relevant differences in visuomotor behavior between primates and rodents, including the frontal versus lateral eye locations, presence or absence of a fovea, and the relative reliance on eye versus combined head and eye movements to change gaze. However, recent work suggests that healthy rodents make both voluntary, saccade-like eye movements and they exhibit resetting saccades (quick phases), similar to humans (Harris and Dunn 2023; Liu et al. 2016; Meyer et al. 2020; Michaiel et al. 2020; Sakatani and Isa 2007; Stahl 2004; Zahler et al. 2021). We assessed the kinematics of saccade-like eye movements made while mice were tracking a moving grating and found the amplitude and velocity of these saccades to be similar between Tau and healthy mice. Our findings contrast with a recent report in tau transgenic zebrafish that show slowing and reduced frequency of horizontal saccades, as well as restricted eye movements and low OKR gain (Bai et al. 2024). The more severe oculomotor deficits in the zebrafish model may be explained by more widespread Tau overexpression and overt cell death; we and others have not observed cell death in brain of the Tau transgenic mouse model (Allen et al. 2002; Xu et al. 2014). All these experiments examined horizontal saccades, while vertical saccades are affected first in PSP (Hardwick et al. 2009; Joshi et al. 2010).

We measured vertical saccades in a dome in which stimuli covered the entire visual field (Harris and Dunn 2023). Human subjects readily make voluntary saccades to a visual target; mice make such saccades relatively infrequently. To address this problem, we took advantage of the fact that voluntary saccades and resetting saccades are thought to share immediate premotor neural mechanisms (Siobhan Garbutt et al. 2003). Moreover, resetting saccades are also impaired in people with PSP (S. Garbutt et al. 2004; Hale et al. 2024). We found that in response to continuous full-field visual stimuli, WT and Tau mice make resetting saccades, consistent with previous observations in healthy mice (Al-Khindi et al. 2022; Harris and Dunn 2023; Yonehara et al. 2016). In Tau mice, we found a selective deficit in the frequency of vertical resetting saccades, as has been observed in PSP (S. Garbutt et al. 2004). However, we did not find slower saccades in Tau mice, in contrast to what is observed early PSP (Shaikh et al. 2017). Saccadic eye movements, especially vertical ones, tend to be small in mice (Sakatani and Isa 2007). It is possible that the low amplitudes of vertical resetting saccades in mice reduced our ability to detect differences in the velocity between WT and Tau mice.

Impaired vertical eye movements in Tau mice establish these mice as a potential model for PSP. While there are differences between mouse, non-human primate, and human oculomotor systems, they share certain fundamentals. For example, much like in primates, mouse extraocular muscles are innervated by projections from motoneuron pools within the oculomotor (III) and abducens (VI) nuclei (Bohlen et al. 2019). We recorded eye movements in a classic head-fixed configuration, easing data analysis and interpretation. However, this configuration doesn’t reflect the typical stimuli eliciting eye movements in mice. Future studies can assess eye and head movements in freely-moving Tau mice as their changes in gaze are commonly mediated by both head and eye movements (Meyer et al. 2018; Meyer et al. 2020; Michaiel et al. 2020). Importantly, measuring the eye movement response to vestibular stimuli should provide useful information to compare with both deficits in visually driven eye movements and the more general motor control deficits in Tau mice. Recording and comparing the movements of both eyes, under both monocular and binocular viewing, in response to optokinetic stimuli might also be helpful in describing the Tau phenotype since subcortical structures strongly influence optokinetic responses under different viewing conditions. In the future, the mouse genetic toolbox can further help dissect the pathological processes underlying the vertical eye movement impairments we found in Tau mice.

Though PSP has classically been associated with pathology in the cerebral cortex and midbrain, a recent comprehensive study of postmortem tissue found early accumulation of Tau pathology in several other motor control regions, including the basal ganglia (Gabor G. Kovacs et al. 2020). We detected Tau pathology in many of the same brain regions in the mouse model, but the relative density of such pathology clearly differs between PSP and the mouse model. Basal ganglia regions such as the SNr, STN, and globus pallidus show high levels of pathology early in PSP (Gabor G. Kovacs et al. 2020), but show relatively sparse pathology in Tau mice. The pattern of Tau pathology is likely due to the fact that, in this transgenic line, human Tau is expressed using the Thy-1 promoter. In the mouse brain, Thy-1 is expressed primarily in neurons and varies between neuronal cell types, with the highest expression levels in cortical projection neurons, and much lower if any expression in basal ganglia regions (Barlow and Huntley 2000; Morris 1985). If anything, this relatively limited Tau expression might have reduced our ability to detect oculomotor phenotypes; in a PSP clinico-pathological correlation study, the burden of Tau pathology in the SNr correlated with severity of gaze palsy (Halliday 2000).

We did, however, find that pathology density in select brain regions correlated with gait impairment, reinforcing the potential importance of Tau pathology in mediating disease phenotypes. More specifically, we found Tau pathology in the PPN correlated with gait and balance impairment. Part of the mesencephalic locomotor region, the PPN has been implicated in gait across multiple species, including humans (Lau et al. 2015) and mice (Caggiano et al. 2018; Dautan et al. 2021; Piallat et al. 2009; Roseberry et al. 2016). Indeed, a recent study found that overexpressing Tau in PPN cholinergic neurons was sufficient to impair coordination in rats (King et al. 2021). Pathology in several other sensorimotor regions also correlated with gait and balance deficits. The inverse correlation between VN pathology and rotarod performance is consistent with the involvement of the vestibular nuclei in balance (McCall et al. 2016; Murray et al. 2018). We looked for pathological correlates of vertical eye movement deficits, and of the regions examined, only DCN pathology showed a strong correlation. It is possible that oligomeric or less aggregated forms of Tau (not detected with our histological assay) are present in regions more classically associated with vertical eye movement control, and that these regions contribute to eye movement deficits.

In sum, Tau mice recapitulate some key features of PSP, opening the door to future mechanistic studies. With this phenotypically penetrant mouse model, interventions such as Tau reduction can be tested for their ability to rescue the impairments in gait and vertical saccades. Moreover, future physiological studies can dissect how Tau pathology affects circuit function to produce these behavioral deficits. Understanding how Tau pathology mediates disease phenotypes at the cellular and circuit level will lead to new therapeutic directions for PSP.

## Materials and Methods

### Animals

Homozygous hTau.P301S transgenic mice on the C57Bl/6 background of both sexes were obtained from Michel Goedert (MRC Laboratory of Molecular Biology, Cambridge, UK) and bred in our colony. Transgenic mice were bred to homozygosity. Age-matched wild-type C57Bl/6 mice were bred in our colony. Mice ranging from 1 to 6 months were used in this study. Animals were housed 1-5 per cage on a 12-hour light/dark cycle with ad libitum access to food and water. All behavioral experiments were performed during the light phase and with approval from the University of California, San Francisco Institutional Animal Care and Use Committee.

### Gross Locomotor and Gait Analysis

Detailed methods for gross locomotor assessment can be found at: dx.doi.org/10.17504/protocols.io.q26g7yo4kgwz/v1. Mice were assessed on the open field and accelerating rotarod tests at 1 – 6 months of age. For open-field assessment, mice were placed in a cylindrical transparent acrylic cylinder and allowed to explore for 20 minutes. Mice were tracked using an overhead camera. Videos were analyzed using Ethovision XT software (Noldus) to obtain the mouse velocity. Average velocity from the last 10 minutes were reported. For the accelerating rotarod (Ugo Basile), mice were placed on the rod, which ramped from 5 to 80 rpm over 300 sec. Latency to fall is the time from the start of the trial to the time when mice either lost grip (rotated with the rod for one revolution) or fell off. Mice performed 3 trials in a session with a 5-minute intertrial interval. The average latency to fall across the 3 sessions was plotted.

Detailed methods for gait assessment can be found at: dx.doi.org/10.17504/protocols.io.n2bvjnwjbgk5/v1. Mice walked back and forth on a linear track (transparent acrylic, 33 inches long, 3 inches wide) between enclosed areas (one opaque black acrylic, one clear acrylic) on either end. This approach yielded 3 to 4 traverses of the track per session. A high-speed camera (60 fps) aimed at a mirror located underneath the track was used to capture the movement of the mouse’s paws. Videos were subsequently tracked using DeepLabCut (see Analysis section).

### Surgery

Detailed surgical methods can be found at: https://www.protocols.io/view/mouse-stereotaxic-surgery-n2bvj6qynlk5/v1. Animals used for eye movement recordings were fitted with a skull-mounted head fixation device. Briefly, anesthesia was induced with intraperitoneal (i.p.) ketamine/xylazine (40/10 mg/kg) and maintained with inhaled isoflurane (1%). In the stereotaxic frame (Kopf instruments), hair was removed and the scalp was cleaned with 3 rounds of alternating Betadine and ethanol prior to making a midline incision. The skull was scraped of overlying membranes, and a thin layer of dental cement (Metabond, Parkell) was applied to the surface of the skull. Mice were instrumented with either a custom head-plate or custom fitted flush press inserts (brass, E-Z Lok 240-000-BR, 0-80). Dental acrylic (light curing composite, Henry Schein) was then used fix the device in place. Mice were then returned to their home cage and administered analgesics (ketoprofen and buprenorphine). Mice were allowed 5-7 days to recover prior to eye movement recordings.

### Eye movement recordings

#### Bidirectional Optokinetic Reflex Stimulus

To measure the slow phase of the OKR along the horizontal plane, we used a custom-designed behavior rig (Liu et al. 2016). Briefly, three LED computer monitors (Dell, 22 inch) were mounted orthogonally to form an enclosure that covered approximately 270 degrees of the visual field along the azimuth (horizontal plane). Visual stimuli were generated with Psychophysics Toolbox 3 using MATLAB (MathWorks). To ensure synchronized movement across our three monitors we used AMD Eyefinity Technology (Radeon Pro WX 5100).

The monitors displayed a vertical sinusoidal grating where the spacing between the stripes was adjusted throughout the horizontal plane such that the spatial frequency of the grating was perceived as constant throughout the visual field, as if the grating was drifting along the surface of the virtual drum. The grating oscillated clockwise and counterclockwise (amplitude: +/-5°, grating spatial frequency: 0.2 cycles/°; oscillation frequency: 0.2, 0.4, 0.6, 0.8, and 1.0 Hz). The stimulus lasted 10 seconds, with 4 seconds of static grating before and after the stimulus (8 second interstimulus interval). Eye movements were recorded with a video camera under infrared illumination at a sample rate of 200Hz.

#### Unidirectional Optokinetic Reflex Stimulus

To measure OKR in both horizontal and vertical directions, and elicit resetting saccades, we used a custom-designed behavior rig providing full-field visual stimuli (Harris and Dunn 2023). A detailed protocol can be found here: dx.doi.org/10.17504/protocols.io.4r3l2qjbjl1y/v1. Briefly, a 24-inch diameter acrylic hemisphere was covered with custom paint with 50% diffuse reflectivity between 350 and 750nm to limit reflections. Stimuli was emitted from a DLP projector and reflected onto the hemisphere via a 6-inch convex mirror.

Here, the visual stimuli were unidirectional gratings presented in six consecutive epochs, consisting of upwards (3 epochs) and downwards (3 epochs) movement for vertical eye movements or leftward (3 epochs) and rightward (3 epochs) for horizontal eye movements. The direction of visual stimuli was consistent within an epoch but randomized across the session. Each epoch consisted of 5 seconds of static grating followed by 30 seconds of grating drifting either directly upward or downward, and an additional 5 seconds of static grating. All gratings moved at a 10°/second and had a spatial frequency of 0.15 cycles/°. Eye movements were recorded with a video camera under infrared illumination at a sample rate of 130Hz.

#### Calibration

The movement of the pupil was calibrated using the corneal reflection generated by an infrared LED affixed to the top of the camera. This point was used as a reference for the eye position. Mice were presented with a static virtual drum and the eye and pupil centers were identified. The camera was then rotated 10 degrees left and right in alternating fashion to calculate the distance between the pupil reference and the pupil center. The dynamic range of pupil size was captured by step-wise increase in the luminance of the static virtual drum.

### Histology and Microscopy

Mice were terminally anesthetized with ketamine/xylazine (200/40 mg/kg i.p), transcardially perfused with 4% paraformaldehyde (PFA) in phosphate buffered saline (PBS), and the brain removed from the skull. Brains were post-fixed for 24 hours in the same PFA solution and then transferred to a solution of 30% sucrose in PBS at 4C. 30-micron sagittal sections were prepared on a freezing microtome. Sections were stained using a standard protocol: https://www.protocols.io/view/immunohistochemistry-14egn7nezv5d/v1. Briefly, sections were washed in 1X PBS 3 x 5 min at room temperature, then blocked for 1 hour in 1X PBS containing 3% normal donkey serum and 0.1% Triton-X. Sections were then incubated in primary antibody cocktail including rabbit anti-tyrosine hydroxylase (1:1000, Pel-Freez Biologicals, cat. #P40101-150), mouse anti-phospho-Tau (AT8, 1:1000; Thermofisher, cat # MN1020) and guinea pig anti-NeuN (1:1000; Millipore cat. # ABN90) for 36-48 hours at 4C. Sections were washed in 1X PBS then transferred to secondary antibody (Jackson ImmunoResearch Labs: 1:500 in 3% NDS in 1X PBS) overnight at 4C. The following day, sections were mounted onto slides and coverslipped using Vectashield mounting medium with DAPI. Low magnification stitched images were obtained using a Zeiss Axio Scan 2.1 microscope at 20X. High magnification images (40X Z-stacks) were obtained using the Olympus Fluoview FV3000.

## Statistical Analysis

Statistical analysis of all experiments is summarized in Table S2 and were performed using GraphPad Prism 10 (GraphPad Software, Inc. San Diego, CA). All data are presented as mean ± SEM. We did not assume normality and used non-parametric statistical tests. Sample size was determined based on estimated effect sizes from published papers using similar mice or methods. Statistical significance was set at a P-value < 0.05.

### Gait analysis

For linear track videos, body parts including the left and right forelimbs, left and right hindlimbs, base of the tail, and nose were tracked with a trained neural network using DeepLabCut. The resulting .csv file containing x,y coordinates for each body part (left and right forelimb, left and right hindlimb, nose, and tail) were then analyzed using custom code (<https://github.com/UCSF-Nelson-Lab/Catwalk-Analysis> ) based on previously described methods (Machado et al. 2015). Periods where the mouse walked in a straight line were manually isolated and the trajectories of the body parts were extracted. Limb trajectories were normalized to the midpoint between tail and nose. Start and end of individual strides were identified by taking the maximum value of the left forelimb trajectory (i.e., when the limb is closest to the nose), corresponding to the initiation of the stance phase. The minimum limb trajectory value (i.e., when the limb is farthest from the nose) corresponds to the end of the stance and start of the swing phase.

For each individual stride we obtained the cadence, stride length, swing fraction, stance fraction, and walking speed. These parameters were compared between genotypes (Fig. 1F, S1C, and Table S1). To calculate stepping variability, the start of the stepping cycle (i.e., start of stance phase) was referenced to the start of the left forelimb, and the stepping time from the other three limbs were calculated as angle with respect to the left forelimb. The standard deviation of these angles was used as a measure of the stepping variability (Fig. 1G, S1D). These were then averaged across limbs and comparisons was made between WT and Tau mice using the Mann Whitney U test (Fig. 1H, S1E).

### Eye movement analysis

Files were imported into MATLAB and analyzed using custom code (<https://github.com/UCSF-Nelson-Lab/OKR-Gain-Analysis>). For OKR gain analysis, trials (∼15-20 trails per temporal frequency) were averaged across the respective temporal frequencies and those containing spontaneous saccades were removed. Gain was calculated as the ratio of the difference of the maximum and minimum of the eye position for each sinusoidal cycle and the amplitude of the linear grating (Eye _amplitude_ / Drum _amplitude_). For detection of horizontal saccades, we used a custom MATLAB function (Zahler et al. 2021) (https://github.com/UCSF-Nelson-Lab/OKR-Gain-Analysis). Events were considered saccades if they had a minimum amplitude of 3° and a minimum velocity of 100°/s. The same function was used to analyze spontaneous and resetting saccades. As vertical saccades are generally smaller, the minimum amplitude was set to 1° to help with detection. The amplitude and peak velocity were obtained from the function.

### Tau pathology quantification

Tau pathology was quantified in sagittal sections using QuPath (open-source software for image analysis, https://qupath.github.io/). Detailed method for quantification can be found at: dx.doi.org/10.17504/protocols.io.36wgqnxwxgk5/v1. Briefly, images are imported into QuPath. Using the ABBA (Aligning Big Brain and Atlases) extension from QuPath, sagittal sections were mapped onto the Allen Brain Atlas slices and scaled to fit the tissue. Borders for each region were defined according to the mapped atlas and adjusted manually for more accuracy when appropriate. 5-7 sections were quantified per hemisphere. Tau+ cell bodies were identified and quantified using custom script (https://github.com/UCSF-Nelson-Lab/Qupath-related-scripts-for-Tau-quantification), and then manually verified for better accuracy. For each mouse, density values for each region were exported to Excel for statistical analysis. Correlations between Tau pathology and behavioral outcomes were made using Pearson correlations.

## Acknowledgements

This work was supported by grants from the National Institutes of Neurological Disorders and Stroke (NINDS) and National Eye Institute (NEI) (K00NS108458 to RBC; R01NS101354 to ABN; R01EY035028 to FAD; F31EY033225 to SCH), the Burroughs Wellcome Fund Postdoctoral Diversity Enrichment Fund (#1022360 to RBC), and an endowment from Richard and Shirley Cahill and Family (ABN). We thank Dr Michel Goedert for providing the hP301S mice. We thank Drs. Massimo Scanziani and Dorit Ron, as well as members of their laboratories for helpful suggestions and use of equipment. We thank all members of the Nelson lab for helpful feedback on the project and manuscript.

## Author contributions

RBC and ABN conceptualized and designed the experiments. DSZ and SdL contributed to experimental design. GB aided in building OKR rig, as well as experimental design, and provided code to extract data for analysis. RBC performed all locomotor, gait/balance, and bidirectional grating experiments and analysis. RBC and SCH performed unidirectional grating experiments and analysis. RBC and SS performed all sectioning, staining, and imaging, and SS performed the QuPath analysis for Tau quantification.

## References

Al-Khindi, T., et al. (2022), ‘The transcription factor Tbx5 regulates direction-selective retinal ganglion cell development and image stabilization’, Curr Biol, 32 (19), 4286-98.e5.

Allen, Bridget, et al. (2002), ‘Abundant Tau Filaments and Nonapoptotic Neurodegeneration in Transgenic Mice Expressing Human P301S Tau Protein’, The Journal of Neuroscience, 22 (21), 9340–51.

Bai, Qing, et al. (2024), ‘A human Tau expressing zebrafish model of progressive supranuclear palsy identifies Brd4 as a regulator of microglial synaptic elimination’, Nature Communications, 15 (1).

Barlow, Jennifer Z. and Huntley, George W. (2000), ‘Developmentally regulated expression of Thy-1 in structures of the mouse sensory-motor system’, The Journal of Comparative Neurology, 421 (2), 215–33.

Baston, Chiara, et al. (2014), ‘Postural strategies assessed with inertial sensors in healthy and parkinsonian subjects’, Gait Ȧ Posture, 40 (1), 70–75.

Bluett, B., et al. (2021), ‘Best Practices in the Clinical Management of Progressive Supranuclear Palsy and Corticobasal Syndrome: A Consensus Statement of the CurePSP Centers of Care’, Front Neurol, 12, 694872.

Bohlen, Martin O., et al. (2019), ‘Mouse Extraocular Muscles and the Musculotopic Organization of Their Innervation’, The Anatomical Record, 302 (10), 1865–85.

Burciu, Roxana G., et al. (2015), ‘Distinct patterns of brain activity in progressive supranuclear palsy and Parkinson’s disease’, Movement Disorders, 30 (9), 1248–58.

Caggiano, V., et al. (2018), ‘Midbrain circuits that set locomotor speed and gait selection’, Nature, 553 (7689), 455–60.

Chen, Athena L., et al. (2010), ‘The Disturbance of Gaze in Progressive Supranuclear Palsy: Implications for Pathogenesis’, Frontiers in Neurology, 1.

Darricau, Morgane, et al. (2023), ‘Tau seeds from patients induce progressive supranuclear palsy pathology and symptoms in primates’, Brain, 146 (6), 2524–34.

Dautan, Daniel, et al. (2021), ‘Modulation of motor behavior by the mesencephalic locomotor region’, Cell Reports, 36 (8), 109594.

De Vos, Maarten, et al. (2020), ‘Discriminating progressive supranuclear palsy from Parkinson’s disease using wearable technology and machine learning’, Gait & Posture, 77, 257–63.

Denk, Franziska and Wade-Martins, Richard (2009), ‘Knock-out and transgenic mouse models of tauopathies’, Neurobiology of Aging, 30 (1), 1–13.

Dutt, Shubir, et al. (2016), ‘Progression of brain atrophy in PSP and CBS over 6 months and 1 year’, Neurology, 87 (19), 2016–25.

Egerton, Thorlene, Williams, David R., and Iansek, Robert (2012), ‘Comparison of gait in progressive supranuclear palsy, Parkinson’s disease and healthy older adults’, BMC Neurology, 12 (1), 116.

Garbutt, S., et al. (2004), ‘Abnormalities of optokinetic nystagmus in progressive supranuclear palsy’, J Neurol Neurosurg Psychiatry, 75 (10), 1386–94.

Garbutt, Siobhan, et al. (2003), ‘Vertical Optokinetic Nystagmus and Saccades in Normal Human Subjects’, Investigative Opthalmology & Visual Science, 44 (9), 3833.

Goedert, M., Jakes, R., and Vanmechelen, E. (1995), ‘Monoclonal antibody AT8 recognises tau protein phosphorylated at both serine 202 and threonine 205’, Neurosci Lett, 189 (3), 167–9.

Guo, Tong, Noble, Wendy, and Hanger, Diane P. (2017), ‘Roles of tau protein in health and disease’, Acta Neuropathologica, 133 (5), 665–704.

Hale, David Edward, Reich, Stephen, and Gold, Dan (2024), ‘Optokinetic nystagmus: six practical uses’, Practical Neurology, 24 (4), 285–88.

Halliday, G. M. (2000), ‘A role for the substantia nigra pars reticulata in the gaze palsy of progressive supranuclear palsy’, Brain, 123 (4), 724–32.

Hardwick, A., et al. (2009), ‘EVOLUTION OF OCULOMOTOR AND CLINICAL FINDINGS IN AUTOPSY-PROVEN RICHARDSON SYNDROME’, Neurology, 73 (24), 2122–24.

Harris, S. C. and Dunn, F. A. (2023), ‘Asymmetric retinal direction tuning predicts optokinetic eye movements across stimulus conditions’, Elife, 12.

Hauw, J.-J., et al. (1994), ‘Preliminary NINDS neuropathologic criteria for Steele-Richardson-Olszewski syndrome (progressive supranuclear palsy)’, Neurology, 44 (11), 2015–15.

Jang, Hochung, et al. (2019), ‘Gait Ignition Failure in JNPL3 Human Tau-mutant Mice’, Experimental Neurobiology, 28 (3), 404–13.

Joshi, Anand C., et al. (2010), ‘Selective defects of visual tracking in progressive supranuclear palsy (PSP): Implications for mechanisms of motion vision’, Vision Research, 50 (8), 761–71.

Kassavetis, Panagiotis, et al. (2022), ‘Eye Movement Disorders in Movement Disorders’, Movement Disorders Clinical Practice, 9 (3), 284–95.

King, Gabriella, et al. (2021), ‘Human wildtype tau expression in cholinergic pedunculopontine tegmental neurons is sufficient to produce PSP-like behavioural deficits and neuropathology’, European Journal of Neuroscience, 54 (10), 7688–709.

Koivisto, Hennariikka, et al. (2019), ‘Progressive age-dependent motor impairment in human tau P301S overexpressing mice’, Behavioural Brain Research, 376, 112158.

Koller, E. J., et al. (2019), ‘Combining P301L and S320F tau variants produces a novel accelerated model of tauopathy’, Hum Mol Genet, 28 (19), 3255–69.

Kovacs, G. G. (2015), ‘Invited review: Neuropathology of tauopathies: principles and practice’, Neuropathology and Applied Neurobiology, 41 (1), 3–23.

Kovacs, Gabor G., et al. (2020), ‘Distribution patterns of tau pathology in progressive supranuclear palsy’, Acta Neuropathologica, 140 (2), 99–119.

Lau, B., et al. (2015), ‘The integrative role of the pedunculopontine nucleus in human gait’, Brain, 138 (Pt 5), 1284–96.

Ling, H. (2016), ‘Clinical Approach to Progressive Supranuclear Palsy’, J Mov Disord, 9 (1), 3–13.

Liu, Bao-Hua, Huberman, Andrew D., and Scanziani, Massimo (2016), ‘Cortico-fugal output from visual cortex promotes plasticity of innate motor behaviour’, Nature, 538 (7625), 383–87.

Machado, Ana S., et al. (2015), ‘A quantitative framework for whole-body coordination reveals specific deficits in freely walking ataxic mice’, eLife, 4, e07892.

Mathis, Alexander, et al. (2018), ‘DeepLabCut: markerless pose estimation of user-defined body parts with deep learning’, Nature Neuroscience, 21 (9), 1281–89.

McCall, Andrew A., et al. (2016), ‘Vestibular nucleus neurons respond to hindlimb movement in the conscious cat’, Journal of Neurophysiology, 116 (4), 1785–94.

Meyer, Arne F., O’Keefe, John, and Poort, Jasper (2020), ‘Two Distinct Types of Eye-Head Coupling in Freely Moving Mice’, Current Biology, 30 (11), 2116-30.e6.

Meyer, Arne F., et al. (2018), ‘A Head-Mounted Camera System Integrates Detailed Behavioral Monitoring with Multichannel Electrophysiology in Freely Moving Mice’, Neuron, 100 (1), 46-60.e7.

Michaiel, Angie M., Abe Elliott Tt, and Niell, Cristopher M. (2020), ‘Dynamics of gaze control during prey capture in freely moving mice’, eLife, 9.

Morris, R. (1985), ‘Thy-1 in developing nervous tissue’, Dev Neurosci, 7 (3), 133–60.

Murray, Andrew J., et al. (2018), ‘Balance Control Mediated by Vestibular Circuits Directing Limb Extension or Antagonist Muscle Co-activation’, Cell Reports, 22 (5), 1325–38.

Narasimhan, Sneha, et al. (2017), ‘Pathological Tau Strains from Human Brains Recapitulate the Diversity of Tauopathies in Nontransgenic Mouse Brain’, The Journal of Neuroscience, 37 (47), 11406–23.

Piallat, B., et al. (2009), ‘Gait is associated with an increase in tonic firing of the sub-cuneiform nucleus neurons’, Neuroscience, 158 (4), 1201–5.

Robinson, John L., et al. (2020), ‘Primary Tau Pathology, Not Copathology, Correlates With Clinical Symptoms in PSP and CBD’, Journal of Neuropathology & Experimental Neurology, 79 (3), 296–304.

Roseberry, T. K., et al. (2016), ‘Cell-Type-Specific Control of Brainstem Locomotor Circuits by Basal Ganglia’, Cell, 164 (3), 526–37.

Sakatani, T. and Isa, T. (2007), ‘Quantitative analysis of spontaneous saccade-like rapid eye movements in C57BL/6 mice’, Neurosci Res, 58 (3), 324–31.

Shaikh Aasef G., Factor Stewart A., and Juncos Jorge L. (2017), ‘Saccades in Progressive Supranuclear Palsy–Maladapted, Irregular, Curved, and Slow’, Movement Disorders Clinical Practice, 4 (5), 671–81.

Stahl, J. S. (2004), ‘Using eye movements to assess brain function in mice’, Vision Res, 44 (28), 3401–10.

Steele, John C., Richardson, J. Clifford, and Olszewski, Jerzy (1964), ‘Progressive Supranuclear Palsy: A Heterogeneous Degeneration Involving the Brain Stem, Basal Ganglia and Cerebellum With Vertical Gaze and Pseudobulbar Palsy, Nuchal Dystonia and Dementia’, Archives of Neurology, 10 (4), 333–59.

Van Alphen, A. M., Stahl, J. S., and De Zeeuw, C. I. (2001), ‘The dynamic characteristics of the mouse horizontal vestibulo-ocular and optokinetic response’, Brain Research, 890 (2), 296–305.

Xu, H., et al. (2014), ‘Memory deficits correlate with tau and spine pathology in P301S MAPT transgenic mice’, Neuropathol Appl Neurobiol, 40 (7), 833–43.

Yonehara, Keisuke, et al. (2016), ‘Congenital Nystagmus Gene FRMD7 Is Necessary for Establishing a Neuronal Circuit Asymmetry for Direction Selectivity’, Neuron, 89 (1), 177–93.

Zahler, Sebastian H., et al. (2021), ‘Superior colliculus drives stimulus-evoked directionally biased saccades and attempted head movements in head-fixed mice’, eLife, 10.

