## Supplemental Figures and Tables for "Tau P301S Transgenic Mice Develop Gait and Eye Movement Impairments That Mimic Progressive Supranuclear Palsy"

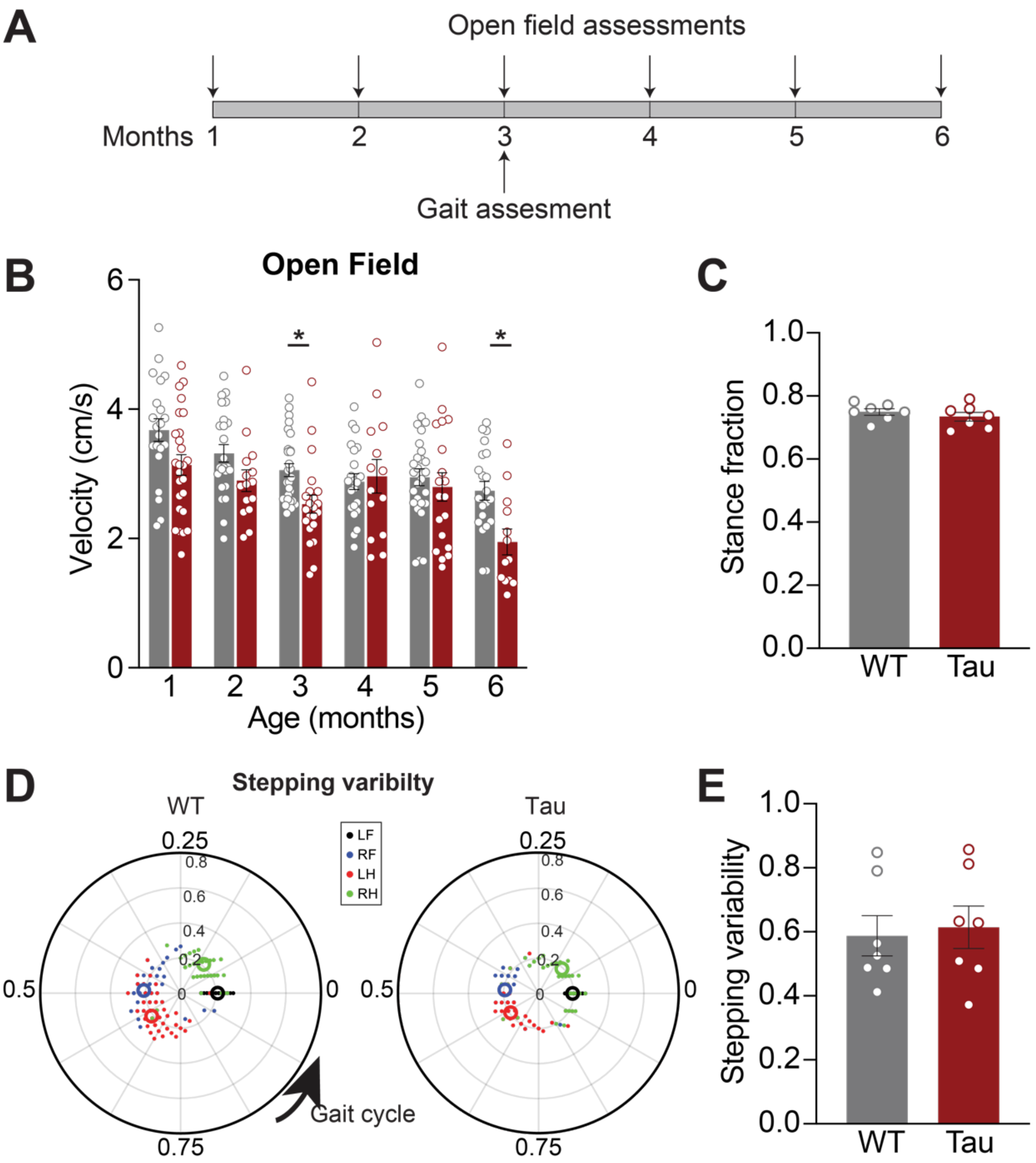


**Figure 1 Supplemental (associated with Figure 1). At 3 months of age, Tau mice do not show impairments in locomotor coordination.**

**A.** Experimental timeline for open field assessment and gait analysis on the linear track. **B.** Average open field locomotor velocity of Tau and WT mice. Tau mice moved more slowly at 3 (N = 28 WT, 22 Tau, p = 0.0246) and 6 months (N = 21 WT, 13 hP301S p = 0.0236). **C.** Average stance fraction in 3-month-old WT and Tau mice (N = 7 WT, 7 Tau, p = 0.5350). **D.** Polar plots indicating phase of the gait cycle where each limb enters stance, aligned to the stance onset of the left forelimb. Individual dots represent a single stride, different colors correspond to different limbs (LF- left forelimb, RF- right forelimb, LH- left hindlimb, RH- right hindlimb). The distance between the center and the dots represents the duration (s) of that stride. Larger open circles represent the average stride for each limb. **E.** Quantification of stepping variability (standard deviation) across limbs in 3-month-old WT and Tau mice (N = 7 WT, 7 Tau, p = 0.7104). N refers to mice. Data is shown as mean +/-SEM. Overlaid open circles in B, C, E represent individual animals.

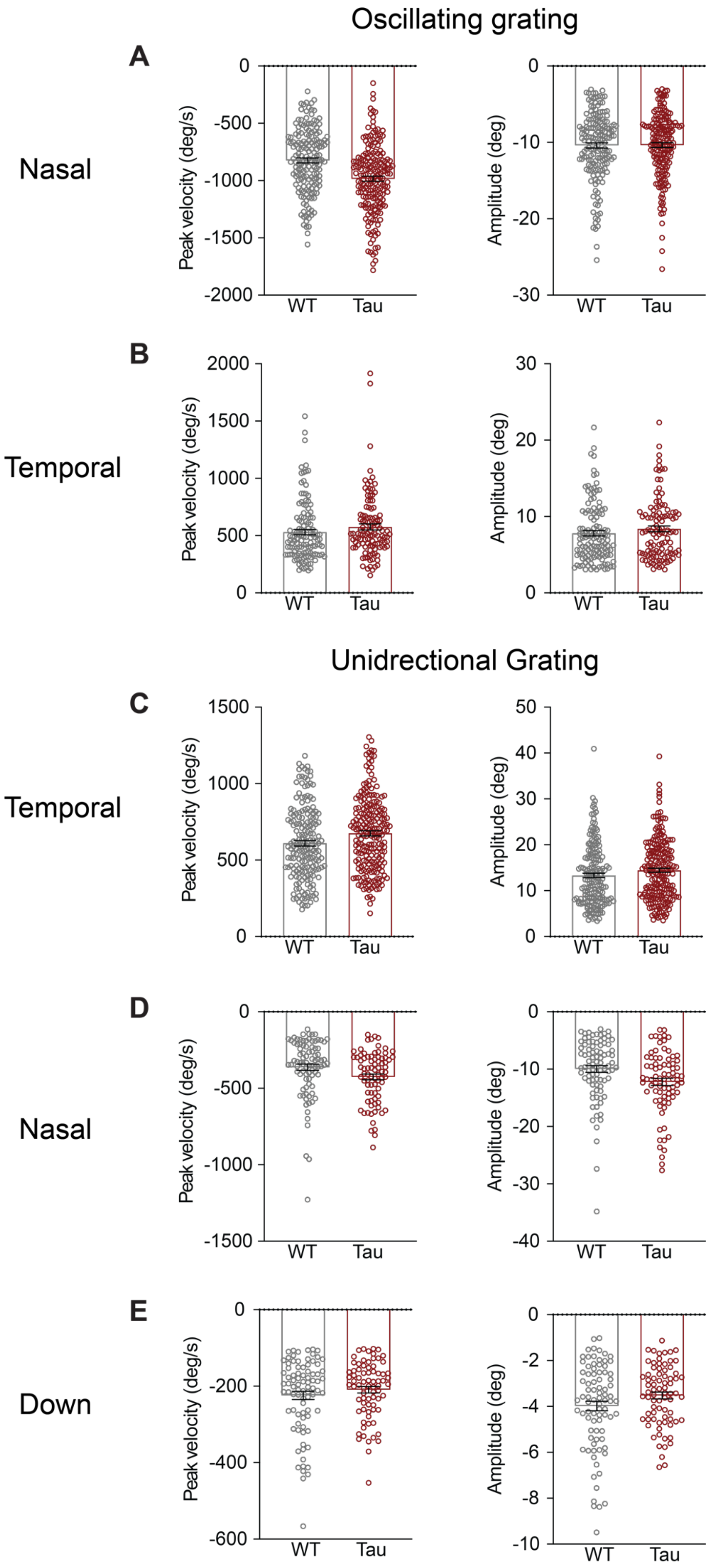
Oscillating grating
A
Nasal
Peak velocity (deg/s)
Amplitude (deg)
WT
Tau
B
Temporal
Peak velocity (deg/s)
Amplitude (deg)
WT
Tau
Unidrectional Grating
C
Temporal
Peak velocity (deg/s)
Amplitude (deg)
WT
Tau
D
Nasal
Peak velocity (deg/s)
Amplitude (deg)
WT
Tau
E
Down
Peak velocity (deg/s)
Amplitude (deg)
WT
Tau

**Figure 2 Supplemental (Associated with figure 3 and 4).**

**A.** Peak velocity (left) and amplitude (right) of nasal spontaneous saccade-like eye movements made during stimulation with a horizontally oscillating grating. WT: N = 9, n = 170, Tau: N = 7, n = 191. **B**. Peak velocity (left) and amplitude (right) of temporal spontaneous saccade-like eye movements made during stimulation with the same oscillating grating WT: N = 9, n = 125, Tau: N = 7, n = 111. **C.** Peak velocity (left) and amplitude (right) of temporal resetting saccades made during stimulation with the horizontal unidirectional grating. WT: N = 8, n = 184, Tau: N = 6, n = 214. **D.** Peak velocity (left) and amplitude (right) of nasal resetting saccades made during stimulation with the horizontal unidirectional grating. WT: N = 8, n= 88, Tau: N = 6, n = 80. **E.** Peak velocity (left) and amplitude (right) of downward resetting saccades made in response to the vertical unidirectional grating. WT: N = 8, n= 84, Tau: N = 6, n = 76. Each open circle represents a single resetting saccade. Data is shown as mean +/- SEM. N = mice, n = saccades.

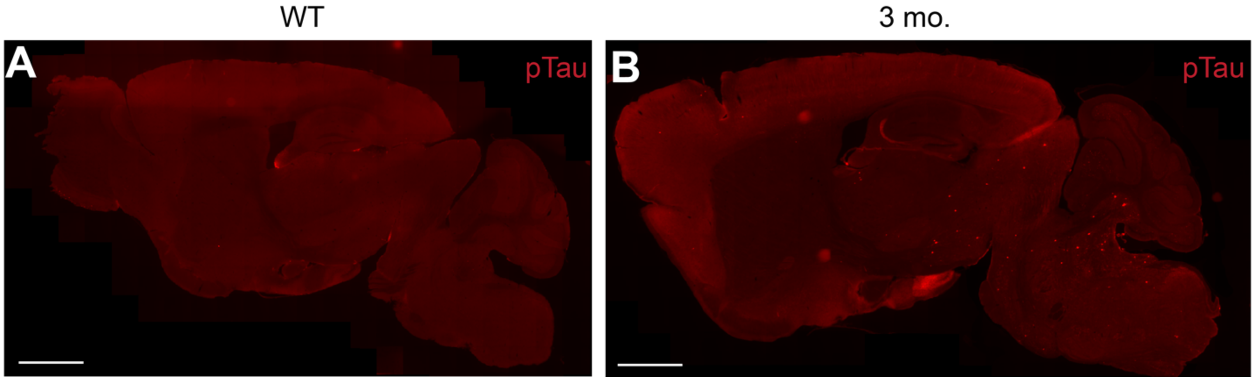


**Supplementary Figure 3 (associated with figure 5). Tau immunoreactivity in WT and 3 mo. old Tau mice.**

**A.** Representative sagittal section from a 5-month-old WT mouse stained for phosphorylated Tau. **B.** Representative sagittal section from a 3-month-old Tau transgenic mouse stained for phosphorylated Tau. Scale bar = 1mm.

**Table S1: GAIT PARAMETERS ANALYSIS AND STATISTICS FOR 5 MO. OLD MICE**
Mann Whitney U test was used for all comparisons.
Table shows statistics for both comparisons using either individual stride or stride averages across mice

| | WT (N = mice, n = strides) | hP301S mice (N = mice, n = strides) | Medians<br>First value- n median<br>Second value () - N median | Comparisons of individual strides | Comparison of individual animals |
|---|---|---|---|---|---|
| Stride length | N = 16, n = 364 | N = 25, n = 707 | WT: 3.678 (3.537)<br>hP301S: 3.171 (3.344) | P < 0.0001 | P = 0.1322 |
| Cadence (steps/ sec) | | | WT: 2.609 (2.640)<br>hP301S: 2.857 (2.849) | P < 0.0001 | P= 0.0434 |
| Speed (cm/s) | | | WT: 8.102 (8.094)<br>hP301S: 8.241 (8.748) | P = 0.5854 | P = 0.7613 |
| Stace fraction | | | WT: 0.6667 (0.6734)<br>hP301S: 0.7273 (0.7401) | P < 0.0001 | P = 0.0013 (Fig. 1E) |
| Swing fraction | | | WT: 0.3810 (0.3670)<br>hP301S: 0.3182 (0.3117) | P < 0.0001 | P = 0.0018 |

**Table S2: Correlations of Tau pathology density to locomotor and oculomotor impairments in 5-6 mo. old Tau mice.**
Spearman correlation was used for all brain regions
Bonferroni Corrected p-values:
Rotarod and Stepping variability – $p < 0.006$
Vertical quick phases – $p < 0.01$

| Brain Region | Rotarod (latency to fall) | Stepping variability (STD) | Vertical quick phase frequency |
|---|---|---|---|
| Motor Cortex | r = -0.7939<br>p = 0.0088 | r = 0.5779<br>p = 0.0525 | N/A |
| Subthalamic Nucleus | r = -0.0424<br>p = 0.9184 | r = 0.4476<br>p = 0.1474 | N/A |
| Zona Incerta | r = -0.6848<br>p = 0.0347 | r = 0.6643<br>p = 0.0219 | r = 0.5798<br>p = 0.2444 |
| Substantia Nigra pars reticulata | r = -0.3212<br>p = 0.3679 | r = 0.5105<br>p = 0.0936 | r = 0.3189<br>p = 0.5444 |
| Pedunculopontine Nucleus | r = -0.7333<br>p = 0.0202 | r = 0.6993<br>p = 0.0142 | N/A |
| Cuneiform Nucleus | r = -0.5273<br>p = 0.1231 | r = 0.3566<br>p = 0.2560 | N/A |
| Superior Colliculus (medial) | N/A | N/A | r = 0.1160<br>p = 0.8444 |
| Medial Vestibular Nucleus | r = -0.9152<br>p = 0.0005 | r = 0.4406<br>p = 0.1542 | r = 0.5444<br>p = 0.3189 |
| Deep Cerebellar Nuclei | r = -0.6606<br>p = 0.0438 | r = 0.6364<br>p = 0.0299 | r = 0.8697<br>p = 0.0333 |

| **Table S3. EXPERIMENTAL ANALYSIS AND STATISTICS**<br>MWU- Mann- Whitney U test; RM-ANOVA- repeated measures analysis of variance | | | | | | |
|---|---|---|---|---|---|---|
| **Key Experiments** | **Figure** | **Comparison** | **Statistical test** | **N (animals)** | **n (strides or events)** | ***p* value** |
| Longitudinal Rotarod analysis | Fig. 1B | Between group | Mixed-effects analysis (Sidak's multiple comparison's test) | WT:<br>1 mo. – 30<br>2 mo. – 30<br>3 mo. – 28<br>4 mo. – 20<br>5 mo. – 21<br>6 mo. – 22<br><br>hP301S:<br>1 mo. – 38<br>2 mo. – 27<br>3 mo. – 22<br>4 mo. – 14<br>5 mo. – 13<br>6 mo. – 15 | N/A | 1 mo. – $p > 0.9$<br>2 mo. – $p > 0.5$<br>3 mo. – $p > 0.1$<br>4 mo. – $p > 0.9$<br>5 mo. – $p > 0.9$<br>6 mo. – $p < 0.0001$ |
| 5 mo. Stance Fraction | Fig. 1F | Between group | MWU | WT = 16<br>hP301S = 25 | N/A | $p = 0.0013$ |
| 5 mo. Stepping Variability | Fig. 1H | Between group | MWU | WT = 16<br>hP301S = 25 | N/A | $p = 0.0002$ |
| Longitudinal open field analysis | Fig. S1B | Between group | Mixed-effects analysis (Sidak's multiple comparison) | WT:<br>1 mo. – 21<br>2 mo. – 24<br>3 mo. – 28<br>4 mo. – 22<br>5 mo. – 27<br>6 mo. – 21<br><br>hP301S:<br>1 mo. – 26<br>2 mo. – 15<br>3 mo. – 22<br>4 mo. – 14<br>5 mo. – 19<br>6 mo. – 13 | N/A | 1 mo. – $p > 0.1$<br>2 mo. – $p > 0.3$<br>3 mo. – $p = 0.0247$<br>4 mo. – $p > 0.9$<br>5 mo. – $p > 0.9$<br>6 mo. – $p = 0.0236$ |
| 3 mo. Stance Fraction | Fig. S1C | Between group | MWU | WT = 7<br>hP301S = 7 | N/A | $p = 0.5350$ |
| 3 mo. Stepping Variability | Fig. S1E | Between group | MWU | WT = 7<br>hP301S = 7 | N/A | $p = 0.7104$ |
| OKR gain | Fig. 2D | Between group | Mixed-effects analysis (w/ Sidak's multiple comparisons) | WT = 9<br>hP301S = 7 | N/A | 0.2: $p = 0.4183$<br>0.4: $p = 0.6412$<br>0.6: $p = 0.9955$<br>0.8: $p = 0.9970$<br>1.0: $p = 0.9750$ |
| Oscillating drum: Nasal main sequence | Fig. 3D | N/A | N/A | WT = 9<br>hP301S = 7 | WT = 170<br>hP301S = 191 | N/A |

| | | | | | | |
|---|---|---|---|---|---|---|
| Oscillating drum: Temporal Main sequence | Fig. 3E | N/A | N/A | WT = 9<br>hP301S = 7 | WT = 111<br>hP301S = 125 | N/A |
| OKR Gain | Fig. 4D | Between group | Two-way RM ANOVA (with Bonferroni correction) | WT = 8<br>hP301S = 6 | N/A | T: p = 0.5185<br>N: p > 0.9999<br>V: p = 0.4430<br>D: p = 0.8834 |
| Linear drum: Temporal main sequence | Fig. 4E | N/A | N/A | WT = 8<br>hP301S = 6 | WT = 184<br>hP301S = 214 | N/A |
| Linear drum: Nasal main sequence | Fig. 4F | N/A | N/A | WT = 8<br>hP301S = 6 | WT = 88<br>hP301S = 80 | N/A |
| Linear drum: Ventral main sequence | Fig. 4G | N/A | N/A | WT = 8<br>hP301S = 6 | WT = 84<br>hP301S = 76 | N/A |
| Nasal Quick phase | Fig. 4H | Between group | MWU | WT = 8<br>hP301S = 6 | N/A | p = 0.8248 |
| Temporal Quick phase | Fig. 4I | Between group | MWU | WT = 8<br>hP301S = 6 | N/A | p = 0.1678 |
| Ventral Quick phase | Fig. 4J | Between group | MWU | WT = 8<br>hP301S = 6 | N/A | p = 0.0373 |
| Dorsal Quick phase | Fig. 4K | Between group | MWU | WT = 8<br>hP301S = 6 | N/A | p = 0.0077 |
| OKR Gain | Fig. 4D | Between group | Two-way RM ANOVA (with Bonferroni correction) | WT = 8<br>hP301S = 6 | N/A | T: p = 0.5185<br>N: p > 0.9999<br>V: p = 0.4430<br>D: p = 0.8834 |
| Oscillating grating: Nasal | Fig. S2A | Between group | MWU | WT = 8<br>hP301S = 6 | WT = 170<br>hP301S = 191 | Velocity: p = 0.3490<br>Amplitude: p = 0.7546 |
| Oscillating grating: Temporal | Fig. S2B | Between group | MWU | WT = 8<br>hP301S = 6 | WT = 111<br>hP301S = 125 | Velocity: p = 0.9497<br>Amplitude: p = 0.8518 |
| Unidirectional grating: Temporal | Fig. S2C | Between group | MWU | WT = 8<br>hP301S = 6 | WT = 184<br>hP301S = 214 | Velocity: p = 0.9497<br>Amplitude: p = 0.8518 |
| Unidirectional grating: Nasal | Fig. S2D | Between group | MWU | WT = 8<br>hP301S = 6 | WT = 88<br>hP301S = 80 | Velocity: p = 0.4908<br>Amplitude: p = 0.4448 |
| Unidirectional grating: Down | Fig. S2E | Between group | MWU | WT = 8<br>hP301S = 6 | WT = 84<br>hP301S = 76 | Velocity: p = 0.7546<br>Amplitude: p = 0.7546 |
